# Excitability in the p53 network mediates robust signaling with tunable activation thresholds in single cells

**DOI:** 10.1101/068668

**Authors:** Gregor Moenke, Elena Christiano, Ana Finzel, Dhana Friedrich, Hanspeter Herzel, Martin Falcke, Alexander LÖewer

**Affiliations:** Mathematical Cell Physiology, Max Delbrueck Center in the Helmholtz Association, Berlin, Germany; Berlin Institute for Medical Systems Biology, Max Delbrueck Center in the Helmholtz Association, Berlin, Germany; Institute for Theoretical Biology, Charité and Humboldt University, Berlin, Germany; Department of Biology, Technical University Darmstadt, Germany

**Keywords:** p53 dynamics, mathematical modelling, single cell analysis, feedback control, excitability

## Abstract

Cellular signaling systems precisely transmit information in the presence of molecular noise while retaining flexibility to accommodate the needs of individual cells. To understand design principles underlying such versatile signaling, we analyzed the response of the tumor suppressor p53 to varying levels of DNA damage in hundreds of individual cells and observed a switch between distinct signaling modes characterized by isolated pulses and sustained oscillations of p53 accumulation. Guided by dynamic systems theory we show that this requires an excitable network structure comprising positive feedback and provide experimental evidence for its molecular identity. The resulting data-driven model reproduced all features of measured signaling responses and is sufficient to explain their heterogeneity in individual cells. We present evidence that heterogeneity in the levels of the feedback regulator Wip1 sets cell-specific thresholds for p53 activation, providing means to modulate its response through interacting signaling pathways. Our results demonstrate how excitable signaling networks can provide high specificity, sensitivity and robustness while retaining unique possibilities to adjust their function to the physiology of individual cells.

## Introduction

To ensure reliable information processing, cellular signaling systems need to faithfully sense inputs in noisy environments while maintaining the flexibility to adjust their function to different physiologies. A commonly observed strategy to enable robust signal detection is the pulsed activation of signaling pathways in a digital-like response ^1^. To understand how pulsatile dynamics can mediate robust yet versatile signal processing, it is necessary to identify the design principles that enable molecular networks to switch between different dynamic states and the mechanisms that allow modulation of their activity.

A well-known example of a pulsatile signaling pathway in mammalian cells is the tumor suppressor p53. As a central hub of the cellular stress response, p53 maintains genomic integrity in proliferating cells and during tissue homeostasis ^2^. In healthy cells, p53 levels are low due to poly-ubiquitination by the E3-ligase Mdm2 and subsequent proteasomal degradation ^3,4^. Upon stress, p53 is activated by kinases that serve as primary damage sensors. One particularly dangerous insult is DNA damage in the form of double strand breaks (DSB), which may cause genomic rearrangements such as translocations, deletions and chromosome fusions. The primary sensor for DSBs is the PI3K-like kinase ataxia telangiectasia mutated (ATM) ^5^, which gets phosphorylated and activated within minutes after damage induction ^6^. Active ATM then stabilizes p53 by at least two distinct mechanisms: it phosphorylates Mdm2, which induces its auto-ubiquitination and subsequent degradation ^7^, and p53, which interferes with Mdm2 binding ^8,9^. As a consequence, p53 accumulates in the nucleus, where it acts as a transcription factor activating the expression of hundreds of target genes ^10^.

A key feature of the signaling network is that p53 transcriptionally activates its own suppressors Mdm2 and the phosphatase PPM1D/Wip1 ^11^, which directly dephosphorylates ATM as well as many ATM substrates such as p53 itself. These interactions constitute negative feedback loops counteracting the p53 response. Using fluorescent reporters and live-cell microscopy, it was previously established that this network architecture generates, at the single-cell level, pulsatile dynamics of p53 accumulation upon DSB induction ^12,13^. Furthermore, it became apparent that the amount of damage present in the cell is not encoded by the amplitude or width of p53 pulses, but rather by the number of uniform pulses in a given time period. However, there was a high degree of heterogeneity, manifested in broad distributions of pulse numbers even in genetically identical cells treated with equal doses of damaging agents. The temporal pattern of p53 pulses showed substantial variability as well: it ranged from regular sustained oscillations in heavily damaged cells to isolated pulses under basal conditions ^14^. Interestingly, no clear threshold in the number of DSBs needed to elicit a pulse could be identified ^15^. Instead, there were indications that the sensitivity of the p53 system was adjusted according to the state of an individual cells. These observations elicit the question how the same molecular network can generate such diverse dynamic responses and how the transition between isolated p53 pulses and oscillatory dynamics is regulated. Furthermore, we are challenged to understand how the p53 response is affected by cellular heterogeneity and how it is adjusted to the needs of individual cells.

To investigate the design principles underlying dynamic signal processing in the p53 network, we combined quantitative single cell data with an abstracted mathematical model of selected molecular interactions. Previous p53 modeling approaches focused on the negative feedbacks mediated by Mdm2 ^16-18^ and Wip1 ^13,19^. Although it is well known that negative feedback loops can give rise to sustained oscillations ^20^, it is less evident how such a system would generate isolated, tunable pulses. To address this question, we employed methods from dynamical systems theory and identified a network architecture based on positive and negative feedback loops that reproduced experimentally measured p53 dynamics over a wide range of conditions. The corresponding mathematical model revealed sources of cellular heterogeneity and allowed us to predict and validate entry points to modulate the sensitivity of the p53 response.

## Results

### Measured p53 dynamics in single cells provide evidence for oscillatory and non-oscillatory regimes

To analyze p53 dynamics in response to different doses of DNA damage, we used previously generated single cell data obtained from a breast cancer cell line expressing a fluorescent reporter system (Fig. 1A, ^14^). Cells were treated with varying doses of the radiomimetic drug neocarzinostatin (NCS, ^21^) to induce a burst of DNA double strand breaks (DSBs) and followed for 48h post damage by time-lapse live-cell microscopy. P53 dynamics were extracted by computer-aided image analysis to obtain time-resolved trajectories for hundreds of individual cells (Fig. 1A-C). To faithfully analyze p53 dynamics despite the presence of inevitable noise from single cell measurements, we established a peak detection algorithm based on wavelets (Fig. S1).

**Figure 1.**
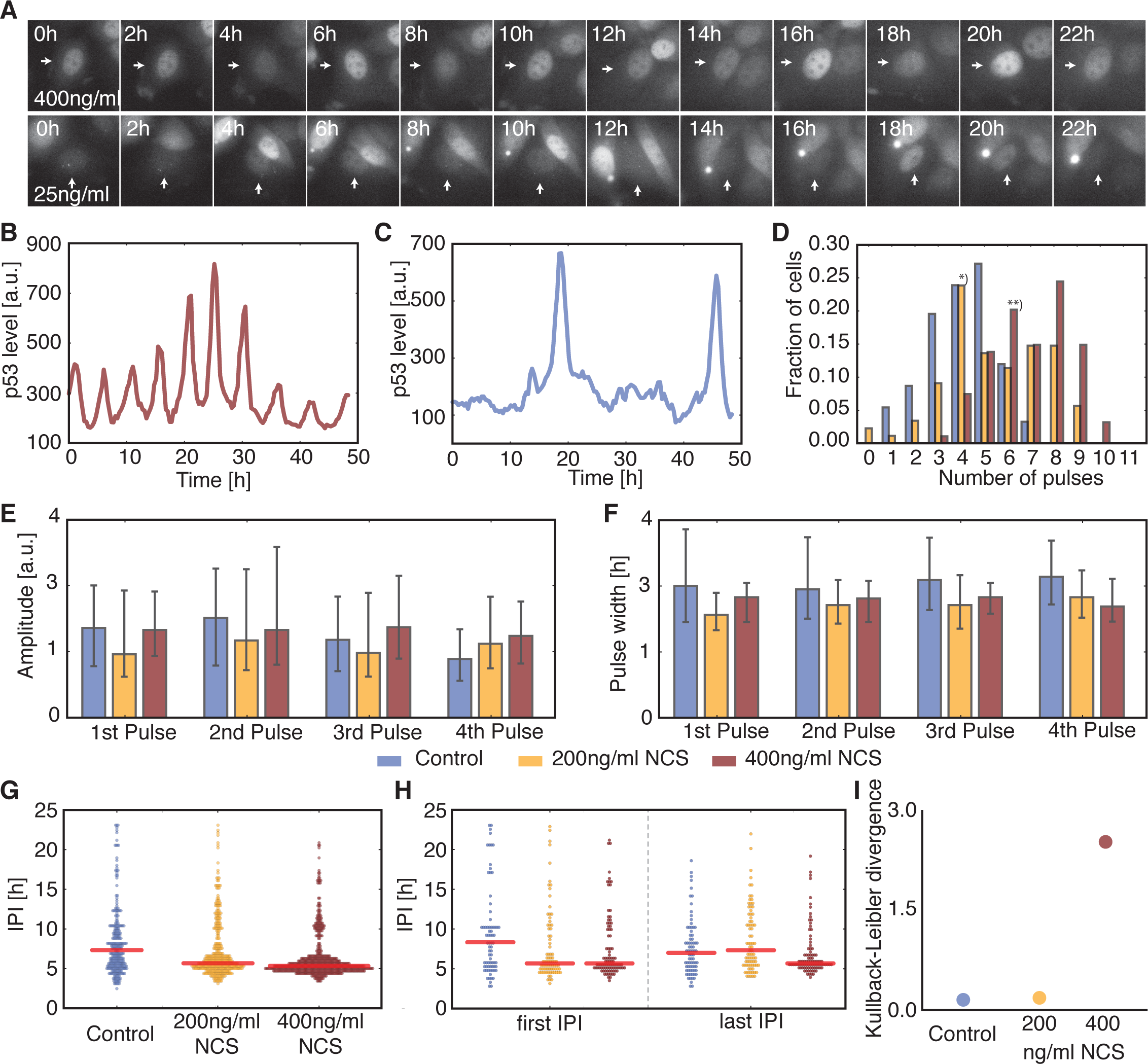
Analysis of p53 single cell trajectories. **A** Time-lapse microscopy images of MCF7 cells expressing p53-Venus after stimulation with 400ng/ml or 25ng/ml NCS. **B-C** Trajectories of p53 protein levels in individual cells showing sustained oscillations (B, upper row in A) or isolated pulses (C, lower row in A). **D** Pulse counting statistics for three different experimental conditions as indicated in the legend below panels E, F. Cells were observed for 48h. **E-F** Median amplitudes (E) and pulse width (F) of the first four p53 pulses, error bars indicate the 1^st^ and 3^rd^ quartile respectively. **G** Inter-pulse-interval (IPI) distributions for all three experimental conditions with samples taken over the entire observation time. Narrower distributions indicate more coherent pulsatile dynamics. **H** IPI distributions of the first and last recorded IPI respectively. The medium stimulated cells (yellow) already returned to irregular pulsatile dynamics at the end of the experiments, while the highly stimulated cells (brown) still show very coherent pulse trains. **I** Kullback-Leibler divergence (KLD) of the first control IPIs from the last IPIs for all three conditions. Dynamics of the medium stimulated cells are indistinguishable from the basal dynamics at later time points.

As previously reported ^12-14,17^, we observed that the number of p53 pulses increased with the damage dose, whereas the pulse amplitudes and widths were on average independent of the stimulation strength (Fig. 1 D, E and F). However, all features analyzed showed a high degree of cell-to-cell variability. To further characterize variability of the pulsatile p53 dynamics we analyzed the distribution of inter-pulse-intervals (IPIs) (Fig. 1G). The spread of an IPI distribution gives a measure for the coherence of a pulsing signal. Oscillatory signals with low noise result in a sharp distribution centered at the mean IPI corresponding to the period of the oscillations. This approach has been previously applied in biology, for example to classify intra-cellular calcium signaling ^22,23^.

When we compared IPI distributions in cells treated with no, intermediate or high levels of NCS, we observed increasing coherence of the p53 pulse trains with increasing stimuli (Fig. 1G). To gain a better understanding of the evolution of p53 dynamics, we analyzed the intervals of the first and last measured pulse separately for each condition (Fig. 1H). In untreated cells, the distributions for the first and last IPIs were similar, indicating that basal p53 dynamics are stationary under our experimental conditions as expected. Upon high levels of DNA damage, we observed narrower distributions centered at the characteristic period of 5h for first and last IPIs, indicating oscillatory dynamics over the entire time period. However, when we analyzed IPIs at medium damage levels, which initially showed the same narrow IPI distribution around 5h, we noticed that the distribution of last IPI was indistinguishable from control cells, indicating that most cells already returned to basal dynamics by that time. We used the Kullback-Leibler divergence, a measure of the distance between probability densities, to support these observations in a quantitative manner. To this end, we defined the distribution of the first IPIs of the control cells as the reference distribution and calculated the divergence between it and the last IPI distributions for all three experimental conditions. The divergence of the control and medium stimulated cells is practically identical, whereas the divergence of the last IPIs of the highly stimulated cells from the first IPIs of the control cells is considerably larger (Fig. 1I).

Our analysis indicates that there are two possible modes of function for the p53 system: Highly stimulated cells show a coherent pulse train resembling oscillations (Fig. 1B) with a sharp IPI distribution centered at a period of about five hours. In contrast, basal dynamics (Fig. 1C) are characterized by isolated pulses without a detectable period, resulting in an IPI distribution consistent with a stochastic process. Cells with intermediate damage alternate between both modes. Importantly, amplitude and width of p53 pulses remain unaffected by these transitions (Fig. 1G-H). These observed p53 dynamics were representative for all cells analyzed, as we did not detect cell death during the time of the experiment.

How can the same molecular network give rise to oscillations and stochastic pulses while preserving pulse amplitude and width? To address this question we turned to dynamical systems theory ^24,25^, whose mathematical analysis allows for the understanding of complex nonlinear phenomena. In the next section we outline and illustrate some key results in the context of the stimulus response of biological regulatory networks.

### Constraining regulatory network topologies by dynamical systems theory

The negative feedback loops by Mdm2 and Wip1 are prominent regulatory elements of the p53 network. It is well known that negative feedback can give rise to limit cycle oscillations ^20^, and most published p53 models take advantage of this to implement pulsatile p53 dynamics ^13,16-18^. To explore whether such negative feedback oscillators are also capable of showing isolated pulses and to examine their behavior when switching between oscillation and steady state, we generated a mathematical model of a simple negative feedback (NF) system (Fig. 2 A and SM section 1). We initialized the system at steady state, mimicking a signal transduction system waiting for input, and applied varying transient stimuli. The system responded with damped oscillations whose maximal amplitudes were dependent on the input strengths (Fig. 2 C). When challenged with a sustained signal, the system showed limit cycle oscillations with fixed amplitude and period (Fig. 2 E, G), resembling the p53 dynamics observed in highly stimulated cells. We again observed a series of pulses with gradually decreasing amplitudes upon decay of the signal.

**Figure 2.**
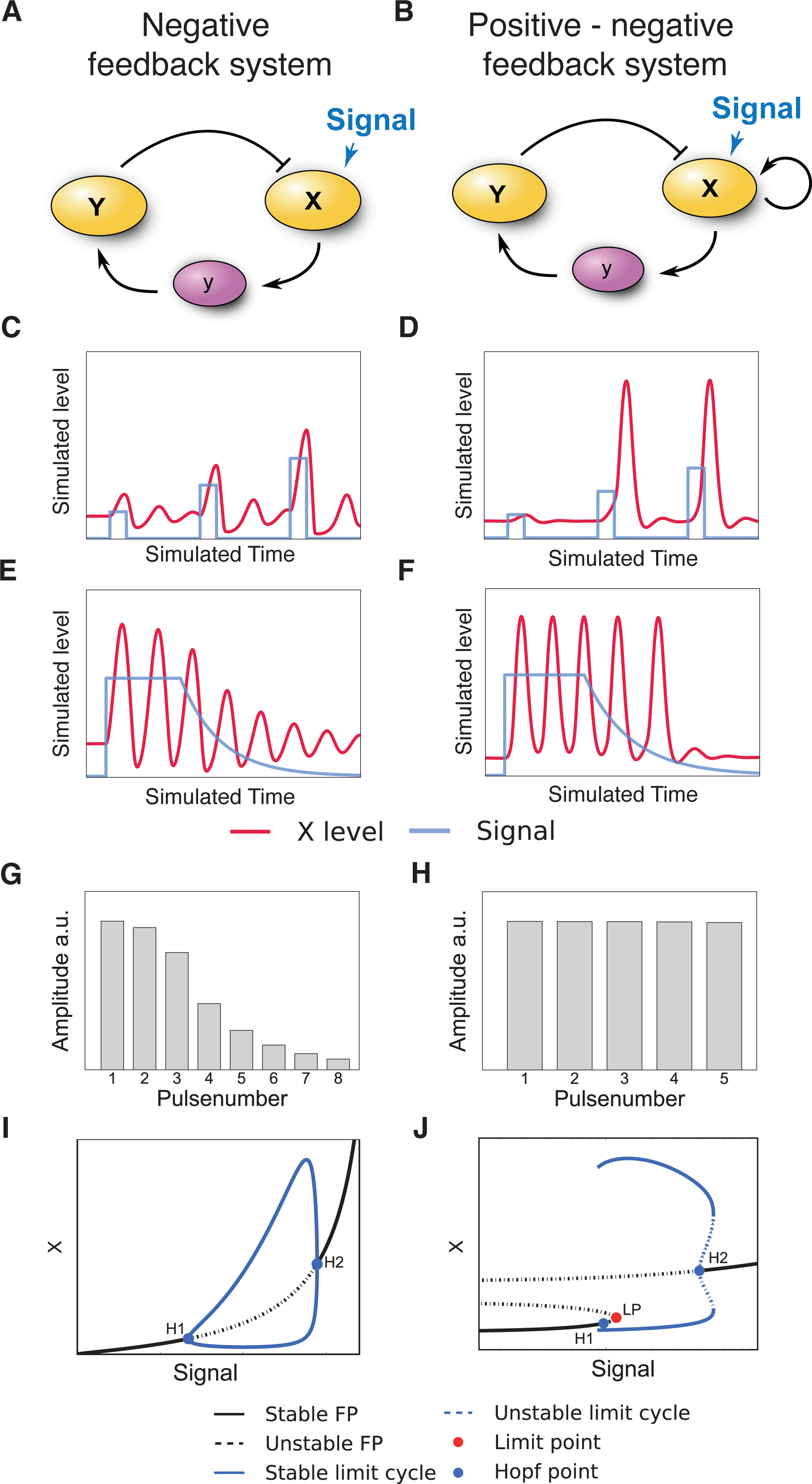
Regulatory network topology constrains dynamic stimulus responses. **A-B** Interaction graphs of hypothetical systems comprising only negative feedback (NF, A) or combined negative and positive feedbacks (NPF, B). The positive feedback is realized by a positive self-interaction. **C-D** Time-resolved response of the NF (C) and NPF (D) systems to transient signals of increasing strengths. **E-H** Time courses demonstrating the termination of oscillations during dynamic switching after sustained stimulation for the NF (E) and NPF (F) system. The corresponding amplitudes of the oscillation were quantified for individual peaks (G and H). Subthreshold responses were excluded. **I** Bifurcation diagram of the NF system for varying signal strengths. The oscillatory regime is confined by two supercritical Hopf bifurcations (H1 and H2). The amplitudes of the limit cycle oscillations are indicated by the blue line. **J** Bifurcation diagram of the NPF system for varying signal strengths. The system is excitable for low input signals. For higher signal strengths a limit cycle is born with large amplitudes around the subcritical Hopf bifurcation (H1). The stable rest state disappears via a saddle-node bifurcation (LP).

To gain a deeper understanding of the dynamic regimes covered by the mathematical negative feedback model, we systematically analyzed its behavior for a range of parameter values. This procedure is known as bifurcation analysis in dynamical systems theory. At low input strength, the system is at a stable steady state. When the signal strength is increased beyond a critical value, the system oscillates. At the critical value, the amplitudes of the oscillations are zero and they increase with increasing signaling strength. This behavior is a generic feature of NF systems. More precisely, systems comprising only negative feedbacks exhibit Hopf bifurcations as the only bifurcation towards an oscillatory dynamic regime (Mallet-Paret and Smith 1990, Pigolotti 2007). As a consequence, NF systems generally show varying amplitudes when switching in and out of the oscillatory regime. However, this is inconsistent with the uniform amplitude distributions observed for p53 pulses in undamaged and damaged cells.

We therefore extended the range of possible p53 network topologies by including a positive feedback (negative-positive feedback (NPF) systems (Fig. 2 B) and supplementary material section 1). In a simple model including positive autoregulation, we observed either no response to a transient stimulation or pulses with amplitudes independent of the stimulation strength for larger stimuli (Fig. 2 D). The existence of such a stimulation threshold separating no response from full response identifies the type of dynamics of our model as excitable ^26^. At stimulation levels below the threshold, negative feedback is stronger than the positive one. Above the threshold value the positive feedback prevails and causes the full response. Importantly, positive feedback is necessary for the existence of this type of excitability ^27^.

The NPF system should also exhibit an oscillatory regime to fully capture the behavior of the p53 system. Indeed, the proximity of excitable and oscillatory regimes with respect to parameter variation is a hallmark of excitable systems. Accordingly, we find for our NPF system that sustained input results in sustained oscillations (Fig. 2F). Importantly, the amplitudes of pulses remain high during signal decay until the oscillations suddenly terminate with only a small subthreshold response detectable (Fig. 2 H).

To better understand the onset of oscillatory dynamics in the NPF system, we again analyzed its behavior upon stimulus variation. There are various types of transitions to oscillations possible for NPF systems, many of which have in common the onset of oscillations with large amplitude ^26^. As shown in the bifurcation diagram (Fig. 2J), oscillations appear with full strength explaining how high amplitude pulses are maintained during signal decay. This abrupt appearance of an oscillatory regime is generic for NPF systems and is unfeasible for NF systems ^24-26^.

Based on these considerations we suggest that the p53 system exhibits two different dynamic regimes controlled by stimulation strength: In basal state the system is excitable and DSB noise causes irregular appearance of pulses by random occurrence of stimulations above threshold. These pulses occur repeatedly in the oscillatory regime at stronger and sustained stimulation, as the input signal is constantly supercritical, explaining the onset of oscillatory pulses with large amplitude.

The dynamics of our abstract NPF model closely resembles p53 behavior observed in basal state, upon intermediate and upon strong stimulation (Fig. 1 H). Excitability at low stimulation and control and the conservation of pulse amplitudes during transitions between dynamic regimes strongly suggest that the p53 network comprises at least one positive feedback.

### Positive feedback on ATM kinase is sufficient to drive excitatory p53 dynamics

As dynamical systems theory suggested the existence of positive feedback in the p53 pathway, we aimed to generate a plausible model of the underlying molecular network. There are numerous positive and negative feedback loops reported for p53, mostly based on the transcriptional activity of the tumor suppressor^28,29^. We monitored RNA levels of candidate feedbacks using qPCR during the first hours post damage (Fig. S2) to determine if these interactions are active in the cell line used to measure p53 dynamics and to test whether they act at a time scale relevant for the pulse formation upon DSB induction. As expected, we did not observe changes in p53 mRNA levels, ruling out positive autoregulation (Fig. S2 A, ^30^). PTEN expression has been reported to be part of a positive feedback loop involving the kinase Akt ^31,32^. Strikingly, we could not detect transcriptional activation of PTEN in MCF7 cells even on longer timescales (Fig. S2 D). We also measured expression levels of the target genes PIDD, reported to induce a positive feedback loop via Caspase-2 mediated Mdm2 cleavage ^33^, and 14-3-3s, which may stabilize p53 by preventing its interaction with Mdm2 ^34^. However, we only observed mild up-regulation of these genes compared to the negative regulator Wip1 or the effector gene p21 (Fig. S2 B-D). Moreover, the expression kinetics of PIDD and 14-3-3s did not match the pulsatile dynamics of p53 accumulation.

As we found no convincing evidence for positive transcriptional feedbacks, we extended our analysis to the upstream kinases that act as initial damage sensors and mediators of the DDR and other feedbacks by phosphorylation and dephosphorylation (Fig. 3A). A central player of DDR initiation is ATM ^5^. In unstressed conditions, ATM forms an inactive homodimer. After the induction of DSBs, ATM gets rapidly activated by a complex formed of Mre11, Rad50 and Nbs1 (MRN), and dissociates into its catalytically active monomeric form upon autophosphorylation ^6^. Activated ATM (ATM^*^) phosphorylates the histone variant H2AX, which serves as a scaffold for the recruitment of further proteins involved in DDR and damage repair. These regions of phosphorylated H2AX (γH2AX) may spread over several thousand basepairs around damage loci ^35^. Subsequently, more MRN is recruited and in turn enhances H2AX phosphorylation and ATM activation ^36^. Although the molecular characterization of the early stages of DDR initiation remains incomplete, it is assumed that this positive feedback enables rapid activation of ATM ^5^ (phosphorylation 1 in Fig. 3A). ATM^*^ subsequently stabilizes p53 by phosphorylating Mdm2 and p53 (phosphorylation 2 and 3 in Fig. 3A, respectively).

**Figure 3.**
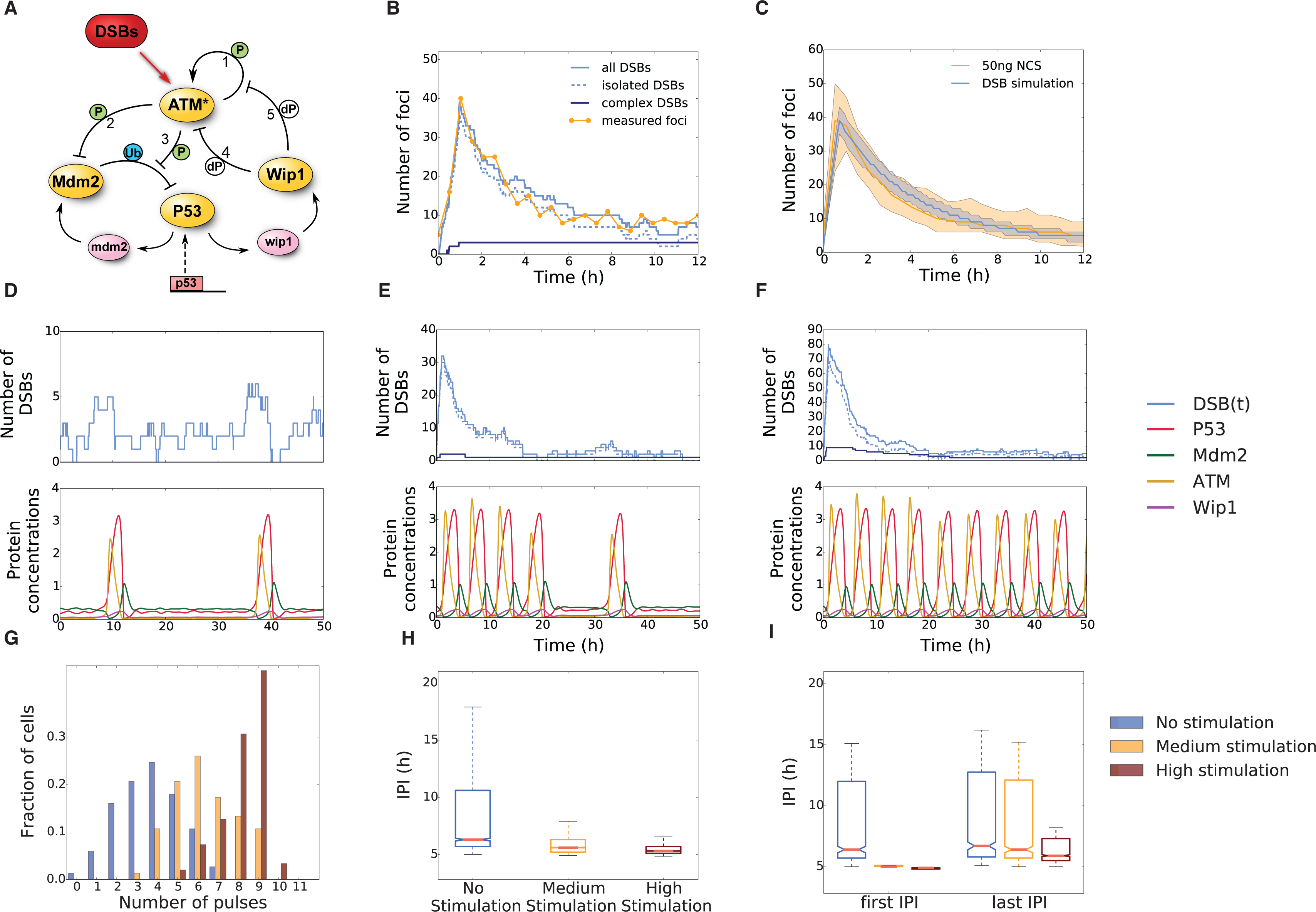
A p53 model driven by stochastic DSB processes recovers experimentally measured dynamical properties. **A** Interaction graph of the modeled p53 network in response to DSBs. Modelled molecular interactions include phosphorylation (P), dephosphorylation (dP) and ubiquitination (Ub); mRNA species (mdm2 and wip1) are shown in purple, all other species describe proteins. The numbers refer to the explanations in the text. **B** A single exemplary realization of the stochastic DSB process. Shown are the dynamics of the two damage types (isolated and complex) and the total effective number of DSBs over time. For comparison, the measured number of DSBs over time post damage is shown for an individual cell (yellow). **C** Statistics of measured and simulated foci for populations of cells. Shown are the median and 1^st^ and 3^rd^ quartiles. N=120 for simulations, N=52 for measurements **D-F** Exemplary DSB process realizations and the corresponding p53 model responses. Shown are three different stimulus intensities: control condition (**D**), medium stimulation (**E**) and high stimulation (**F**). **G** Pulse counting statistics for three simulated ensembles with stimulation strengths as indicated. **H** Inter-pulse-interval (IPI) distributions corresponding to the ensembles shown in **G**. The overall coherence of the pulsatile response increases with stimulation strengths. **I** First and last IPI distributions for the three different stimulation strengths corresponding to the simulated ensembles shown in **G** and **H**. The medium stimulated cells have mostly returned to unstimulated dynamics, whereas the highly stimulated cells still predominantly show coherent oscillations on the single cell level.

However, even if ATM activation is facilitated by a positive feedback, it is not clear *a priori* that this contributes to the excitable p53 response. ATM may only act as an intermediate signal activating an independent excitatory circuit. To distinguish between these possibilities, we again employed the abstract NPF model and either inhibited the activating signal or the species *X* (Fig. S2E). Signal inhibition did not affect the observed ‘all or none’ response of the system: either a full pulse or a subthreshold response were generated. In contrast, inhibition of species *X* severely affected the excitation loop, as it is an essential part of the excitatory circuit. As a consequence, response amplitudes were reduced depending on the time of inhibition (Fig. S2E).

To test whether ATM acts as an activating signal or an essential part of the excitatory circuit, we modulated the activity of the kinase with the inhibitor Wortmannin at different times after damage induction (Fig. S2F and G). We observed a decrease of p53 pulse amplitudes with earlier addition of the inhibitor, indicating an essential role of ATM in generating the excitable p53 response. Our conclusions are further supported by previous experiments in which p53 accumulation was uncoupled from ATM activation ^37^. In this scenario, only damped oscillations were observed, indicating a lack of positive feedback downstream of ATM.

To explore the effect of an ATM mediated positive feedback on the p53 system, we constructed a mathematical model of the network using ordinary differential equations (ODEs) with parameters based on previous publications (Batchelor et al, Fig. 3A). In addition to positive regulation through ATM and negative regulation through Mdm2, the activity of the oncogenic phosphatase Wip1 was critical for the performance of the p53 system. It antagonizes ATM not only by dephosphorylating and inactivating it (dephosphorylation 4 in Fig. 3A), but also by reverting the modification of its substrates including γH2AX, Nbs1 and Mre11 ^38^ (dephosphorylation 5 in Fig. 3A). Albeit the introduction of positive regulation on ATM may at first appear as a minor modification compared to existing p53 models ^39^, it effectively converts a NF into a NPF system and therefore greatly expands the possible range of dynamical responses (Fig. 2).

Due to the prominent role of positive feedback on ATM, we decided to explicitly consider the generation and repair of DSBs in our model. Recently, we measured DSB dynamics in single cells over time and observed considerable variability in their number and half-life, highlighting the stochastic nature of DSB repair ^15,40^. We represent DSB kinetics by a data based modeling approach involving stochastic birth-death processes ^41^. Birth of a DSB is described by a simple zero order reaction with a variable break rate *b*, allowing us to mimic the action of the radiomimetic drug NCS used in live-cell measurements of DSB dynamics (see also supplement). DSB repair was modeled as a first order reaction with a constant repair rate *r*. The population mean number of DSBs is *<DBS(t)> = (N*_*s*_-*N*_*b*_) *exp(-rt) + N*_*b*_, where *t* is the time since stimulation, *N*_*s*_ the dose dependent maximal number of inflicted DSBs and *N*_*b*_ = *b/r* is the *background damage level*. This allows us to estimate the basal break (0.7 DSB/h) and repair rates (0.35 DSB/h) from measured data (Fig. S4 and supplementary material). These rates suggest cells to have on average two DSBs as background damage level even without any external stimulation, which is consistent with previous reports ^14^. The average repair time is given by *<t*_*repair*_*> ~ ln(N*_*s*_*)/r*. Hence, even high numbers of induced DSBs would be repaired after 20 hours with these values of r. Since measurements showed the persistence of DSBs beyond this time, we expanded our model to include DSBs of variable complexity. It has been suggested that complex breaks (cDSBs) occur when several clustered DSBs are generated within a chromatin loop ^42^. In this case, a substantially longer repair time is assumed compared to *isolated DSB* (iDSB), leading to biphasic DSB kinetics ^43,44^. The amount of cDSBs was estimated to be 10% of the overall number of DSBs ^43^. We modified our model accordingly to represent the total number of DSBs as the sum of stochastic processes defined for iDSBs and cDSBs. We assigned a half-life of 20 hours to cDSBs, which is in accordance with experimental measurements ^45^. This approach allowed us to reproduce the dynamics and variability of DSB induction and repair with reasonable precision (Fig. 3 B and C).

To test whether the combination of an excitable p53 model and stochastic DSB kinetics are sufficient to describe single cell dynamics of p53 in different conditions, we simulated three varying levels of DNA damage. Without externally induced DSBs, average background damage did not elicit a p53 response and the system resided mainly at steady state with subthreshold fluctuations (Fig. 3 D). However, small bursts of DSBs occurred due to the stochastic nature of the DSB process and excited single isolated p53 pulses when crossing the activation threshold (Fig. 3 D). When we simulated an ensemble of cells, we observed wide IPI distributions corresponding to non-oscillatory dynamics (Fig3 H and I). Hence, the experimentally observed basal dynamics of p53 were well explained by the excitable regime of our model (Fig. 1E and F, ^14^).

Next, we simulated a medium damage dose through a higher initial break rate, leading to a substantial increase in DSBs (Fig. 3E). The system responded with an initial period of oscillations. When repair was completed, the network passed from the oscillatory to the excitable regime with essentially unaltered pulse shapes (Fig. 3 E). This transition was also reflected by the IPI distributions of simulated ensembles: the first IPIs displayed a very coherent response, whereas the distribution of the last IPIs was indistinguishable from unstimulated cells (Fig. 3 I). This behavior was comparable to the behavior of cells challenged with intermediate levels of DNA damage (Fig. 1 F).

Finally, we simulated a strong damage dose resulting in high amount of DSBs including many cDSBs. As a result, DNA damage was not entirely repaired within the simulated time; the p53 system received sustained input throughout the simulation and showed oscillatory dynamics (Fig. 3 D and I). In simulated ensembles, we observed coherent IPI distributions for the first and last pulse. The pulse number, however, was still variable, as observed in living cells (Fig. 3 G and Fig. 1 D).

In summary, our model with excitable and oscillatory dynamics was able to reproduce the main characteristics of p53 behavior in single cells: a smooth transition between steady state, irregular pulses and regular oscillations while preserving pulse amplitudes and durations. This is based on the combination of negative and positive feedbacks, which lead to a bifurcation scheme (Fig. S3 A) similar to the one obtained for the simple NPF model (Fig. 2 J). Importantly sensitivity analysis showed that our modeling results were robust against parameter variations (Fig. S4).

### Sensitivity of the p53 response is modulated by Wip1 levels

So far, the only source of variability was given by the stochastic DSB process as the deterministic core model was identical for all cells. Hence, the same initial DSB time course would lead to a collective first response, either a pulse or no pulse, within the population. However, it was previously demonstrated that in clonal cell populations the number of responsive cells gradually increases with the amount of induced DSBs ^15^, suggesting that each individual cell has a different sensitivity for DNA damage. As we can directly correlate the cellular responsiveness to the stimulation threshold in our simulations, we employed our excitable p53 model to understand this heterogeneity. We first investigated how random short-lived fluctuations in protein concentrations influence the stimulation threshold in the excitable regime. To this end, we perturbed the system from steady state and observed whether it generated a full pulse or decayed back with subthreshold dynamics. Combining numerous random perturbations, we could identify the position and orientation of the excitation threshold in phase space (Fig. 4 A). While the position of the threshold was hardly affected by p53 and Mdm2 concentrations, higher levels of Wip1 strongly increased the amount of active ATM needed to trigger an excitation loop. This striking dependence of the threshold position on Wip1 levels suggested that the phosphatase might be a major determinant for a cells individual responsiveness towards DSBs. The special importance of Wip1 for the model performance is also reflected in the sensitivity analysis of the model: parameters associated with Wip1 show the highest sensitivity (Fig. S4).

**Figure 4.**
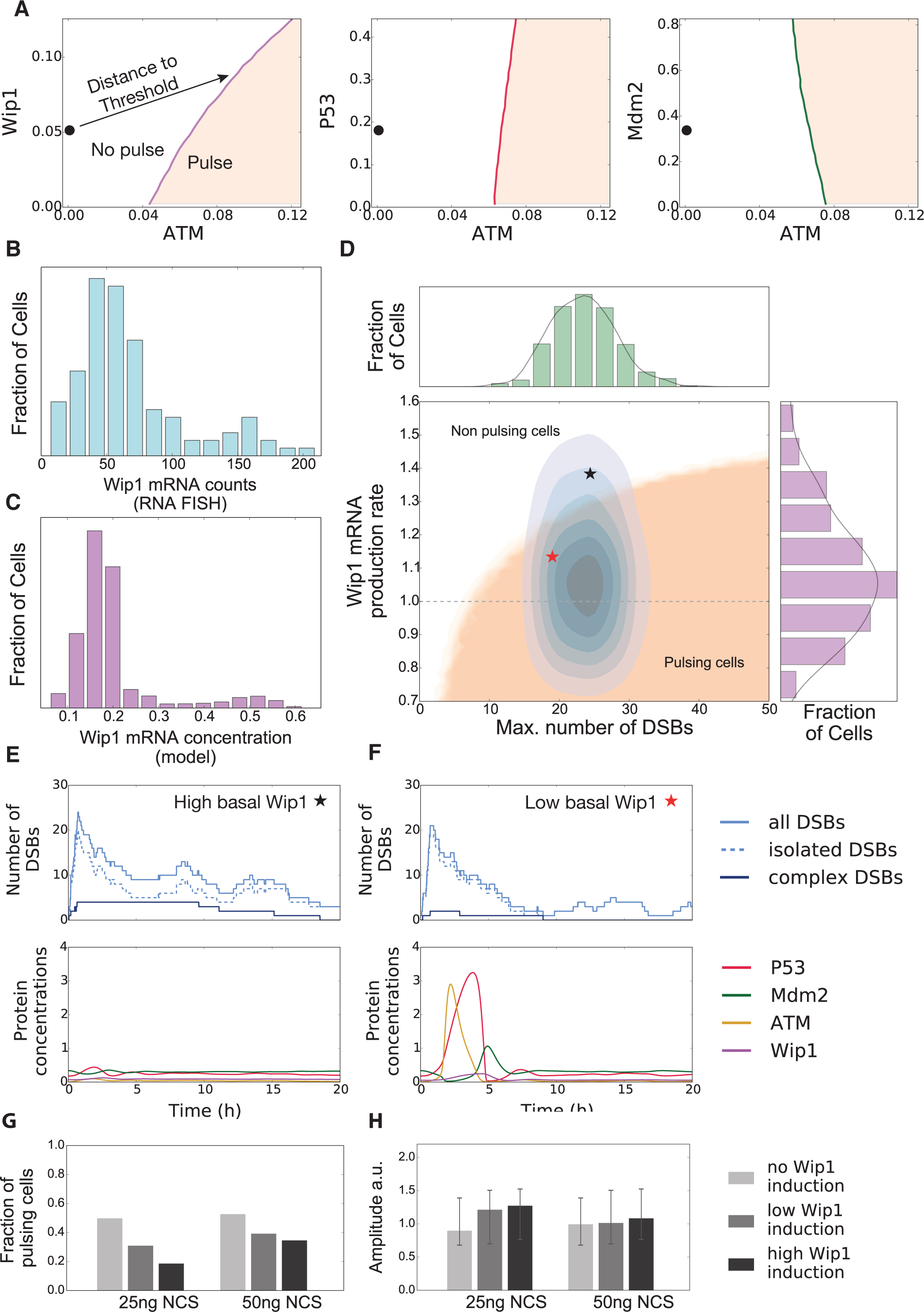
The responsiveness of individual cells towards damage is modulated by Wip1 levels. **A** Sensitivity of the excitation threshold for fluctuations of the three regulated proteins p53, Mdm2 and Wip1. The steady state is depicted as (•) for basal activated ATM levels on the left respectively. The p53 system is particularly sensitive to Wip1 fluctuations, as it influences the threshold the most. **B** Measured single cell Wip1 mRNA counts for an unstimulated cell population (N = 106). **C** Modeled *in silico* Wip1 mRNA concentration variability. The single cell expression levels were drawn from a log-normal distribution and the system was driven by basal DSB dynamics. **D-F** The dependency of p53 responsiveness on damage levels (maximal number of inflicted DSBs) and Wip1 expression rate is shown in D. Cells responding with a p53 pulse lie inside the beige area. A typical fixed dose experiment will induce a distribution of cells inside this plane (blue contours), as both the number of DSBs and the Wip1 expression levels are variable on the single cell level. The fraction of cells within the beige area determines the population response. Exemplary simulations of cells with high (E, black star) or low (F, red star) Wip1 are shown. Although both cells suffered comparable numbers of DSBs, only the cell with low Wip1 level was responsive. **G** Measured fraction of cells showing a pulse in the first six hours for two different damage doses and three different levels of Wip1 overexpression as indicated. Higher Wip1 levels reduced responsiveness *in vivo*. **H** Pulse amplitude distributions of the responding cells shown in **G**.

Our qPCR analysis (Fig. S2) suggests that Wip1 is predominantly regulated on the transcriptional level. We therefore measured the distribution of Wip1 mRNA in single cells and indeed found strong variability (CV = 0.63) in Wip1 mRNA abundance (Fig. 4 B). To introduce variable Wip1 expression in the model, we assumed a lognormal distribution of Wip1 mRNA production rates, which translates to log-normally distributed Wip1 mRNA and protein concentrations at steady state (Fig. S5 A, see Supplementary Information) (Feinermann 2008, Sorger 2009). The simulated distribution (CV = 0.52) corresponded well to the measured distribution of Wip1 mRNAs in single cells (Fig. 4 C). In addition, the weakly bimodal shape of the measured distribution could be naturally recovered in the model due to cells exhibiting a spontaneous p53 pulse at the moment the population snapshot was taken.

We next simulated thousands of trajectories, each with its individual Wip1 production rate according to the distribution described above and DNA damage represented by the stochastic DSB process. We binned the ensemble according to the maximal number of inflicted DSBs and tested for a pulse within the first six hours. This defines two regions in the DSB-Wip1 production rate plane (Fig 4 D) corresponding to either responsive or unresponsive cells. Our analysis showed that with increasing Wip1 levels, more DSBs were needed to trigger a p53 pulse. In addition to the contribution by varying Wip1 levels, we observed uncertainty introduced by stochastic repair, as some cells repaired DSBs before the excitation threshold was crossed. This led to a graded boundary along the two regions, where cells only trigger pulses with a certain probability.

We next simulated an experiment with a given dose of a damaging agent by first sampling individual Wip1 productions rates and then simulated DSB dynamics with a fixed initial break rate. This results in a two dimensional distribution of cells within the DSB-Wip1 production rate plane (Fig. 4 D, histograms and blue contours). The proportion of cells residing in the pulsing and non-pulsing region determines the population response. This is exemplified by two individual simulations with a similar number of DSB, where only one shows a full p53 pulse due to differences in Wip1 levels (Fig. 4 E and F, red and black star in Fig. 4 D). Taken together, our model simulations suggest that Wip1 levels set a cell-specific activation threshold for the p53 system. To test the model hypothesis that the responsiveness of the p53 system is determined by Wip1 steady state concentrations, we established a cell line overexpressing Wip1 fused to the red-fluorescent protein mCherry. Upon inducing a low dose of damage, we observed that cells with higher levels of Wip1 showed less frequent p53 responses (Fig. 4 G). Strikingly, we did not detect substantial changes in pulse amplitudes upon Wip1 induction (Fig. 4 H). Taken together, this strongly supports the idea that the p53 system functions as an *all-or-none* pulse generator and that Wip1 is a major regulator of its cell-specific excitation threshold towards DSBs.

## Discussion

In this study, we combined insights from dynamical systems theory with experimental measurements to identify design principles that enable robust yet versatile signal processing by the tumor suppressor p53. We showed that an excitable network structure comprised of both negative and positive feedbacks is capable of reproducing the p53 response in healthy and stressed cells. While pure negative feedback systems also generate limit cycle oscillations corresponding to highly damaged cells, they fail to reproduce two key features of p53 dynamics observed in single cells: pulses with uniform amplitudes during entry and exit from the oscillatory regime and the occurrence of isolated pulses triggered by spurious endogenous DNA damage. However, our theoretical considerations only pointed towards the existence of positive feedback without constraining its molecular manifestation. We therefore investigated various postulated transcriptional feedbacks including autoregulation of p53 RNA. However, we did not observe expression changes at the level and timescale expected for regulators of pulse formation (Fig. S2). While we cannot formally exclude contributions from additional transcriptional feedbacks, we provide evidence that the switch-like activation of ATM upon damage induction ^6^ followed by rapid degradation of Mdm2 ^39^ provide sufficient feedback upstream of p53. As details of the molecular interactions mediating the initial DNA damage response are still emerging, we decided to capture its structure in an phenomenological model condensing molecular details such as ATM autophosphorylation and its interaction with the MRN complex, MDC1 and phosphorylated H2AX at break sites into a single autoregulatory interaction ^5,36^. Importantly, there are reports of other molecular species like USP10 ^46^ or micro RNAs like miRNA29 ^47^ which support p53 activation, and further experimental and theoretical studies will be needed to decipher the role of individual molecular interactions.

Combining positive feedback around ATM with a regulatory module based on negative feedback regulation resulted in stable p53 levels in healthy cells with few DSBs. Only when a threshold for ATM activity was reached, the system responded with a single full amplitude pulse. This *all-or-nothing* response is characteristic for excitable systems such as the well-known FitzHugh-Nagumo model ^48^. Excitable systems can be classified into two categories ^26^, where type I excitability has the distinct feature of a direction dependent threshold. Such systems are insensitive to perturbations not associated with the primary input. In our p53 model, a pulse can only be triggered by changes in the activity of ATM, the primary sensor for DSBs, while the system is inert to fluctuations in other species. This direction dependent threshold leads to a strong coupling of pathway activation to a specific input. In addition, the *all-or-nothing* response characteristic of excitable systems allows for high sensitivity as the positive feedback essentially serves as a signal amplifier. Therefore, type I excitable systems naturally provide specificity and sensitivity to enable robust signal processing. The information content of a single pulse of defined amplitude is necessarily binary, suggesting that type I excitable systems are well suited to transmit decisive signals. Accordingly, it has been described that the stochastic entry into and exit from the competent state of the bacterium *Bacillus subtilis* is regulated by such a system ^49^.

The p53 system, however, does not only signal the presence or absence of DSBs, but also encodes the extent of DNA damage through the number of uniform pulses in a given time period ^12-14,17^. This is achieved by a seamless transition between incoherent pulsing and sustained oscillations. Sustained input above the activation threshold shifts the system in parameter space from the excitable to the neighboring oscillatory regime. The time spend in this regime effectively determines the number of pulses generated in a given time interval. To analyze signal encoding, we coupled our excitable p53 model to a stochastic process describing the induction and repair of DNA damage, which was parameterized using single-cell measurements of DSB kinetics. This allowed us to reproduce single cell trajectories resembling cells with varying degrees of DNA damage. Stochasticity introduced through DSB kinetics was sufficient to also reflect experimentally observed heterogeneity in pulse numbers and allowed us to reproduce corresponding distributions in population of cells. Variable initial numbers and half-lives of DSBs kept each cell an individual amount of time within the oscillatory regime. The sooner cells returned to the excitable regime, the broader the distribution of inter-pulse intervals became, resulting in increasing heterogeneity in pulse numbers. Taken together, observed cell-to-cell variability in pulse numbers and inter-pulse-intervals can mainly be attributed to a highly stochastic DNA double strand break induction and repair coupled to a type I excitable systems.

In addition to heterogeneity in pulse numbers, single cell measurement also revealed cell-specific activation thresholds for the p53 response ^15^. We therefore analyzed the influence of all model species on the stimulation threshold and found that its value was mainly determined by the phosphatase Wip1. This can be mechanistically understood by taking into account that Wip1 not only directly dephosphorylates ATM, but also interferes with ATM self-activation, e.g. by dephosphorylating γH2AX and the MRN complex ^38^. Our analysis predicted that cell-to-cell variability in Wip1 protein abundances strongly modulates the cellular responsiveness towards DSBs. This is supported by pronounced heterogeneity in Wip1 mRNA levels measured in individual cells. Moreover, we were able to modulate p53 responsiveness on the single cell level by over-expressing Wip1. Our results highlight that excitable systems can provide unique possibilities to combine robust *all-or-nothing* responses with versatile regulation of pathway sensitivity by threshold repositioning.

As a prominent modulator of the p53 system, Wip1 provides an entry point for crosstalk between the DNA damage response and other major signaling pathways. It has been reported that its expression levels are regulated by MAP kinase signaling ^50^, NF-κB activity ^51^ and the tumor suppressor HIPK2 ^52^. This allows to efficiently sensitize or attenuate p53 responsiveness depending on the state of the cell or information from the surrounding tissue and may be important when cells rapidly proliferate during development and regeneration. It also provides a mechanism for inactivating p53 function during tumorigenesis, which is often initiated by aberrant signaling. The oncogenic role of Wip1/PPM1D is highlighted by the frequent observation of gene amplification or activating mutations in human cancers ^53-56^.

There is growing evidence that pulsatile intracellular dynamics play an essential role in different cellular contexts ^57^. They can be observed in distinct biological processes such as signaling through calcium ^23^, MAPK ^58^ or NF-κb ^59,60^, the circadian clock ^61^ or embryonic patterning ^62^. Are there general design principles for the structure of cellular pulse generators? In this study, we use abstract mathematical modeling and experimental measurements to emphasize the importance of the feedback and bifurcation structure for understanding the dynamics and function of a regulatory circuit ^63^. We have found positive feedback and excitability to provide robustness towards cell variability for p53 signaling and previously for Ca^2+^ spiking ^23^. In this scenario, noise and fluctuations play central roles, as they elicit pulses and spikes in the excitable regime. In the future, similar approaches may enable identification of other common principles underlying biological oscillators and may open new avenues to modulate critical cellular processes in the context of human diseases.

## Material and Methods

### Live-cell Imaging

The MCF7 p53 reporter cell line has been described before ^14^. To generate cells with increased PPM1D/Wip1 levels, a pRRL-based lentiviral construct expressing a fusion between Wip1 and mCherry under the control of the constitutive EF1alpha promoter was cloned using the MultiSite Gateway Three-fragment system (LifeTechnologies). Stable clonal cell lines were established following viral infection and selection with puromycin. Cells were maintained at 37 °C / 5% CO_2_ in RPMI 1640 containing 10% FCS, penicillin / streptomycin and appropriate selective antibiotics (400 µg/ml G418, 50µg/ml hygromycin and 0.5µg/ml puromycin) to maintain transgene expression.

For imaging, we seeded cells in poly-D-lysine-coated glass-bottom plates (MatTek Corporation) two days before experiments. The day of the experiment, media was replaced with fresh one lacking phenol red and riboflavin. Cells were imaged on a Nikon Ti inverted fluorescence microscope with a Hamamatsu Orca R2 camera and a 20x plan apo objective (NA 0.75) using appropriate filter sets (Venus: 500/20 nm excitation (EX), 515 nm dichroic beam splitter (BS), 535/30 nm emission (EM); CFP: 436/20 nm EM, 455 nm BS, 480/40 nm EX ; mCherry: 560/40 nm EM; 585 nm BS; 630/75 nm EM, Chroma). The microscope was enclosed with an incubation chamber to maintain constant temperature (37°C), CO_2_ concentration (5%), and humidity. Cells were imaged every 15-20 minutes for the duration of the experiment using Nikon Elements software.

We used custom-written Matlab (MathWorks) scripts based on code developed by the Alon lab (Cohen *et al*., 2008) and the CellProfiler project (Carpenter *et al*., 2006) to track cells throughout the duration of the experiment. Briefly, we applied flat field correction and background subtraction to raw images and segmented individual nuclei from nuclear marker images using adaptive thresholding and seeded watershed algorithms. We then assigned segmented cells to corresponding cells in following images using a greedy match algorithm.

### Time series analysis

The single cell p53 trajectories were analyzed using wavelet transformations and smoothing splines (Fig S1B). Peak detection was done by identification and subsequent filtering of ridge-lines in the wavelet transform, a strategy first devised for mass spectra analysis ^64^. After p53 pulse times were determined, localized smoothing splines were fitted to a trajectory around every individual pulse (Fig S1A) to reliably obtain pulse amplitudes and widths. All analysis steps were compiled into a custom written Python program using the open source scientific tools package SciPy ^65^.

### Simulations

The deterministic p53 model was formulated as a system of coupled ordinary differential equations (ODEs); the equations are given explicitly in the Supplement. Bifurcation analysis and numerical solutions were obtained with the PyDSTool framework (Clewly R 2007), an open source analysis environment for dynamical systems. The stochastic DSB process was simulated based on Gillespie’s stochastic simulation algorithm (SSA, Gillespie 2002). To address the non-stationary damage kinetics of the NCS stimulation, the break rate was approximated as a step function defined by:

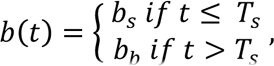

where b_s_ is the break rate during stimulation and b_b_ the basal break rate. The switching time between these two rates is given by T_s_, which corresponds to the time span of active NCS. This approach towards time inhomogeneous Markov processes via step functions still yields analytically tractable expressions for the next-jump densities needed for the SSA algorithm. Details of the implementation and parameter values used can be found in the supplement. Simulations of the stochastically driven p53 system were carried out by pre-computing realizations of the DSB process and augmenting the r.h.s. of the ODE system by a corresponding forcing term.

### qPCR of potential feedback candidates

We extracted mRNA using High Pure RNA Isolation kits (Roche) and generated complementary DNA M-MuLV reverse transcriptase (NEB) and oligo-dT primers. Quantitative PCR was performed in triplicates using SYBR Green reagent (Roche) on a StepOnePlus PCR machine (Applied Biosystems). Primers sequences: β-actin forward, GGC ACC CAG CAC AAT GAA GAT CAA; β-actin reverse, TAG AAG CAT TTG CGG TGG ACG ATG; Wip1 forward, ATA AGC CAG AAC TTC CCA AGG; Wip1 reverse, TGG TCA ATA ACT GTG CTC CTT C; p21 forward, TGG ACC TGT CAC TGT CTT GT; p21 reverse, TCC TGT GGG CGG ATT AG; p53 forward, TGA CTG TAC CAC CAT CCA CTA; p53 reverse, AAA CAC GCA CCT CAA AGC; PIDD forward, GAT GTT CGA GGG CGA AGA G; PIDD reverse, CAG GTG CGA GTA GAA GAC AAA G; PTEN forward, AAG GGA CGA ACT GGT GTA ATG; PTEN reverse, GCC TCT GAC TGG GAA TAG TTA C; TIGAR forward, CCT TAC CAG CCA CTC TGA GC; TIGAR reverse, CCA TGT GCA ATC CAG AGA TG; 14-3-3σ forward, CCC TGA ACT TTT CCG TCT TCC; 14-3-3σ reverse, GGT GCT GTC TTT GTA GGA GTC

### smFISH

MCF7 cells were cultured for 24h on uncoated coverslips (thickness: #1). Cells were washed and fixed with 2% Paraformaldehyde for 10 min at room temperature, washed again and permeabilized over night with 70% Ethanol at 4°C. Custom probe sets for single molecule FISH ^66^ labeled with CalFluor-610 were designed using *Stellaris RNA FISH probe designer* (Biosearch Technologies) on the reference sequence NM_003620.3 (PPM1D). Hybridization was performed at a final concentration of 0.1 µM probe following manufacturers instructions (*Stellaris RNA FISH Protocols-adherent cells*). Coverslips were mounted on Prolong Gold Antifade (Life technologies). For single molecule RNA quantification, 11 z-stacks of each cell were acquired with 300 nm step-width. Quantification of RNA counts per cell was performed using the *Star Search* analysis tool for spot detection (http://rajlab.seas.upenn.edu/StarSearch/launch.html).

## Acknowledgements

We thank Andrea Grybowsky for technical assistance and all members of the Falcke and Loewer labs for helpful discussions. G.M. and A.L. were supported by the German Research Foundation (G.M.: research training group “Computational Systems Biology”; A.L.: priority program “InKoMBio”).

## Author Contribution

G.M. performed data analysis and mathematical modeling; D.F. performed smFISH and A.F. qPCR measurements; E.C. contributed important reagents; G.M. and A.L. prepared figures and wrote the manuscript; H.H., M.F. and A.L. conceived the study and supervised the research.

